# Modular barcode beads for microfluidic single cell genomics

**DOI:** 10.1101/2020.09.10.292326

**Authors:** Cyrille L. Delley, Adam R. Abate

**Affiliations:** Bioengineering and Therapeutic Sciences, University of California, San Francisco, San Francisco, CA 94158, USA; California Institute for Quantitative Biosciences, University of California San Francisco, San Francisco, CA 94158, USA; Chan Zuckerberg Biohub, San Francisco, CA 94158, USA

## Abstract

Barcode beads allow efficient nucleic acid tagging in single cell genomics. Current barcode designs, however, are fabricated with a particular application in mind. Repurposing to novel targets, or altering to add additional targets as information is obtained is possible but the result is suboptimal. Here, we describe a modular framework that simplifies generation of multifunctional beads and allows their easy extension to new targets.

**One Sentence Summary:** We describe an optimized design for barcoding beads which are useful for single cell RNA and DNA sequencing.

## Introduction

Biological samples from tissues or the environment often contain communities of distinct cell types. Such mixtures can be deconvoluted *a priori* by isolating subsets for analysis, or *a posteriori* by employing methods which provide measurements at single cell resolution. For example, single cell genomics (scDNAseq) allows identification of the clonal composition of cancer cells, while single cell transcriptomics (scRNAseq) enables deconvolution of mixed cells with distinct phenotypes(*1*). Because single cell approaches require little prior knowledge about the sample and because recent microfluidic methods provide enough throughput to characterize large and heterogeneous cell populations, these methods are becoming ubiquitous in biological research.

A key step in single cell genomics workflows is loading droplets with high concentrations of barcode oligonucleotides to tag nucleic acids of interest; the resultant tagged molecules can be pooled for all cells and efficiently sequenced in a single run. Barcode loading is often accomplished using beads on which barcode sequences are synthesized(*2, 3*) which provides two advantages: Efficient microfluidic techniques allow most cells to be paired with a bead(*4*) (Figure 1a), while efficient synthesis of oligos on beads provides ample barcode for target tagging, yielding optimal sequencing data. However, current barcode designs utilize fixed targeting primers that are not easily repurposed; consequently, if new targets are identified, a new batch of beads must be synthesized, which is expensive and laborious. For example, repurposing beads used for scRNAseq to target genomic DNA results in barcodes containing poly-T stretches that prevent common sequencers from reading into downstream sequences; nor can additional targets be easily added to existing whole transcriptome or multiplexed amplicon beads, which would allow, for instance, sensitive capture of guide RNAs used in genome wide knockout screens(*5, 6*). If a universal barcode bead could be designed that could be easily and cost effectively retargeted to new sequences, it would allow easy repurposing of existing beads to new targets, accelerating single cell experiments and reducing cost and waste.

**Figure 1.**
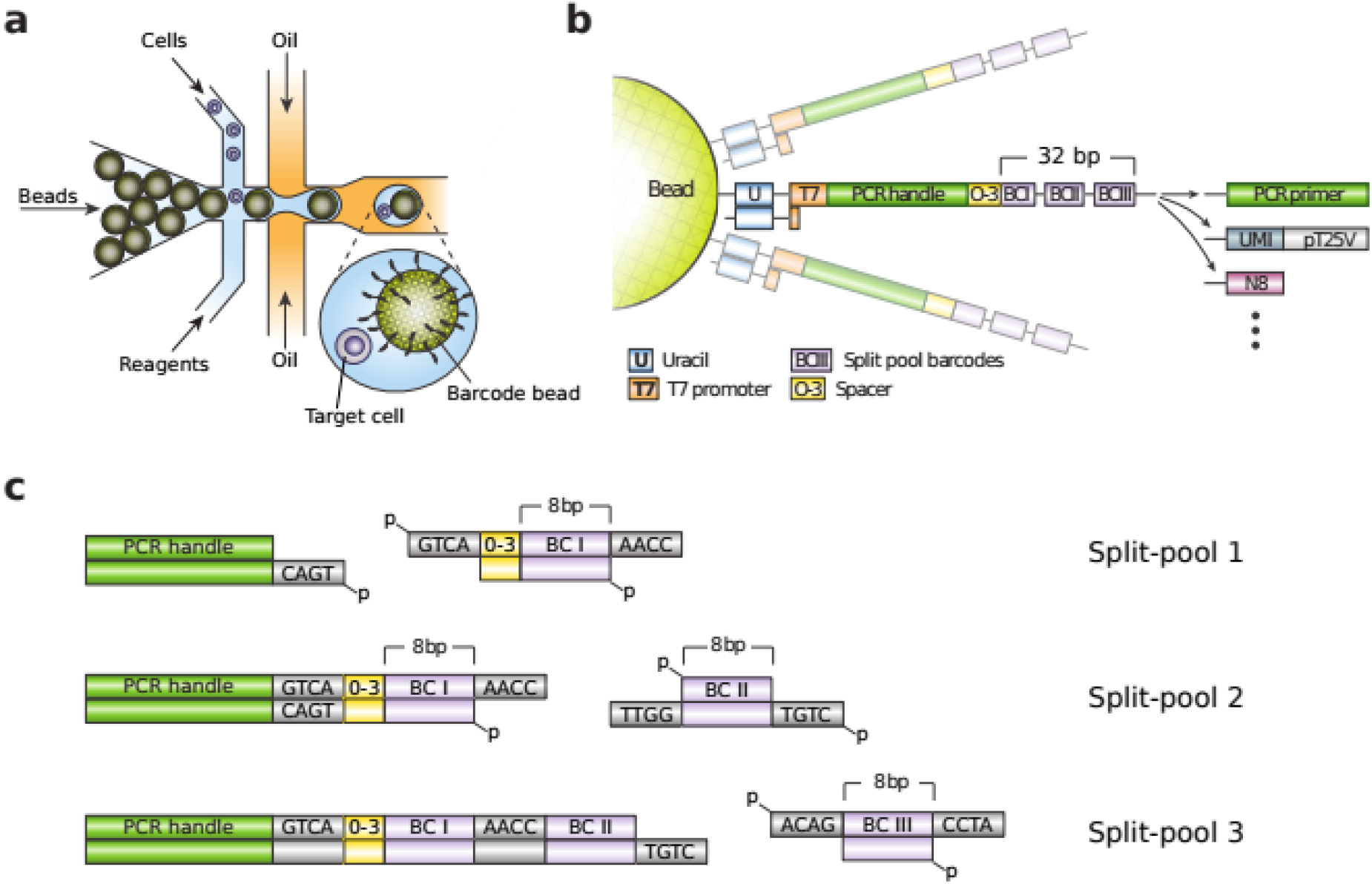
Barcode bead fabrication overview: **a** Barcode beads form the backbone of microfluidic single cell genomic protocols. Close packing enables efficient bead loading. **b** Organization of the barcode sequence on the beads. The elements are modular and can be removed or replaced with other sequence blocks if desired. The 0-3 bp spacer introduces a barcode specific frame shift that helps with cluster detection in Illumina’s sequence technolgoy. **c** DNA ligation with T4-ligase is used to minimize the footprint of the split-pool barcode fragments enabling more split pool cycles per length.

Here, we describe a modular framework that allows easy repurposing of existing bead batches to new targets (Figure 1b, 1c)(supplementary protocol). The modular design is compatible with described single cell sequencing workflows, including scDNAseq(*7*), scRNAseq(*2*), Abseq(*8*), multimodal analyses(*9–12*), and genome wide knockout screens(*13–15*). Moreover, the optimized structure generates compact barcodes that can be assembled in a fraction of the time and cost compared to existing methods. The platform is general and flexible, making it useful for numerous single cell genomics applications.

## Results

### Compact barcode synthesis by split-pool ligation

The core concept of our bead design is to use modular sequence assembly in the synthesis process. Unlike existing approaches which synthesize and target beads in one workflow, thereby yielding beads limited to one sequencing task, we design beads in which the barcode sequence is assembled in a first step, and appropriate primers as targeting moieties are added in a second step, such as poly-T for scRNAseq or multiplexed panels for scDNAseq.

Barcode bead fabrication starts with microfluidic emulsification of a gel precursor solution serving as the bead scaffold(*16, 17*). Once polymerized, we recover the beads and the covalently linked primers serve as anchors for barcode synthesis.

To synthesize barcode sequences on the beads, we use a split-pool approach. Split-pool assembly works by repeatedly partitioning the beads into random fractions, modifying the beads with a specific barcode fragment and pooling the beads. This results in a barcode set which grows exponentially with the number of cycles. Because the acrylamide hydrogel backbone of the beads is incompatible with phosphoramidites oligo synthesis we use enzymatic reactions to concatenate pre-synthesized oligos. As concatenated subsequences we use octamers from a library with minimum Levenshtein distance four(*18, 19*), which enables robust barcode identification even with sequencing errors and indels from oligo synthesis (supplementary data table 1-6).

DNA polymerases have previously been used in split-pool protocols, but require handles for specific hybridization, inflating barcode length. Barcodes can also be assembled with DNA ligases which can operate with four or less base pair overhangs(*20–24*) (Supplementary figure 2a, 2b). These overhangs ensure specific ligation, since different sequences used in sequential steps prevent improper propagation of reactive stubs remaining from failed ligations in previous rounds. To characterize the process, we measure ligation efficiency in a split pool synthesis reaction, observing >80% of stubs are ligated per round, such that after three rounds 64% of oligos on the bead are full length (Supplementary figure 2c). In contrast, beads fabricated using polymerases achieve just 36% yield after two rounds(*17*). Thus, our results demonstrate that ligation is more efficient for barcode bead fabrication in split-pool protocols than polymerase extension while also yielding more compact barcodes that reduce sequencing waste.

Compact barcodes allow more sequence diversity in the same oligo length; this is important because barcode diversity sets barcode collision rate and thus limits the number of cells that can be sequenced. For example, ligation permits assembly of three blocks into a barcode while polymerase extension allows only two. The result is that with three 96-well plates of barcode subsequences, ligation encodes 884,736 different sequences, allowing 45,382 cells per experiment for a 5% clash rate. By contrast, using polymerase extension to assemble two-block barcodes from eight 96 well plates yields just 147,456 barcodes, allowing just 7,564 cells to be profiled. To match ligation, polymerase extension would require eighteen 96-well plates.

In terms of cost, ligation is equivalent to polymerase extension per fabricated bead volume. However, the massive reduction in required barcode subsequences (3 rather than 18) affords a significant lower upfront investment. Ligation is also faster and less laborious: It uses double stranded DNA while polymerases require single stranded, obviating the need to denature after each cycle and reducing the number of wash steps. Moreover, the considerably smaller number of oligos makes manual pipetting feasible, whereas synthesis of two-block polymerase libraries from 18 plates requires robotics to ensure quality beads.

### Enzymatic barcode release is cost effective

Popular protocols for single cell genomics release barcodes from beads to increase availability for reverse transcription or PCR priming. This is normally achieved using UV cleavable chemical moieties, like 2-nitrobenzyl, or disulfides that can be broken with a reducing agent(*2, 25*). While fast to cleave, these linkers are expensive (Supplementary table 1) and require care to avoid premature cleavage during bead fabrication and handling. Our protocol instead employs enzymatic cleavage, yielding significant cost savings while also making the approach less sensitive to premature cleavage. Because genomic DNA is a valuable substrate in many single cell experiments, we avoid restriction enzymes including rare cutting Type IIS enzymes. Thus, we incorporate deoxyribose uracil (dU) as the linker, which does not exist in genomic DNA and can be cleaved by an enzyme mix comprising uracil DNA glycosylase and endonuclease III. Oligos containing dU are readily synthesized and inexpensive, and the linker consumes just one base. Moreover, dU containing oligos are stable and specifically cleavable by these enzymes, which are also readily available and inexpensive.

To characterize the efficiency of this barcode release strategy, we incubate beads functionalized with oligos containing dU in PCR and reverse transcription buffer, mimicking conditions of single cell genomics protocols. To measure cleavage efficiency, we use FAM-labelled probes complimentary to the cleavable oligo and measure fluorescence for cleaved beads and positive and negative controls. We observe little activity on single stranded oligos, but near complete cleavage of double stranded in both buffer conditions (Supplementary figure 2d-f). This makes sense because the enzymes natural substrate is double stranded DNA. Thus, to make the single stranded barcodes cleavable, we include an oligo complementary to the cleavage region in the mix, making it locally double stranded and achieving efficient cleavage. These results show that uracil cleavage is an efficient mechanism for barcode release, while reducing bead fabrication cost to about a third (Supplementary table 1).

### Flexible bead usage for scDNAseq and scRNAseq

To demonstrate the benefit of the modular bead design in typical use cases we create a mock cancer cell system consisting of two human cell lines, Raji and K562, and profile their clonal relationship. We select a panel of 49 genomic locations (supplementary data table 8) that are known hot spots for mutations in acute myeloid leukemia, functionalizing the beads with the corresponding set of 49 forward primers, and running the associated microfluidic workflow(*7*). After cell encapsulation, lysis, and chromatin digestion, we merge the droplets with barcode beads, PCR reagents, and enzymes to cleave the uracil linkers, thereby releasing the barcode primers into solution for targeted amplification of the 49 loci (Figure 2a). The resultant sequence data yields high quality reads across the panel for 1,020 cells, with a median of 3447 reads and 45 detected amplicon loci per cell (Supplementary figure 3a, 3b). After aligning the fragments to the human reference genome and performing variant calling, we use hierarchical clustering to assign genotypes via Ward’s minimum variance method(*26*), obtaining two separated clusters (Figure 2b). A comparison of the detected variant calls in single cells with the genotype obtained from a bulk measurement of Raji and K562 cells shows that the clusters indeed represent the two input lines (Figure 2c)(supplementary data table 9). We apply the same workflow to other cell lines such as P493-6, LAX7R; and profile CEM and K562 with the same beads but a different microfluidic strategy(*27*), to demonstrate that the method is insensitive to cellular or microfluidic differences (Supplementary figure 3c-f and Supplementary figure 4a, 4b). These experiments show that our barcode beads enable high throughput single cell genome sequencing with data quality equivalent to traditional beads.

**Figure 2.**
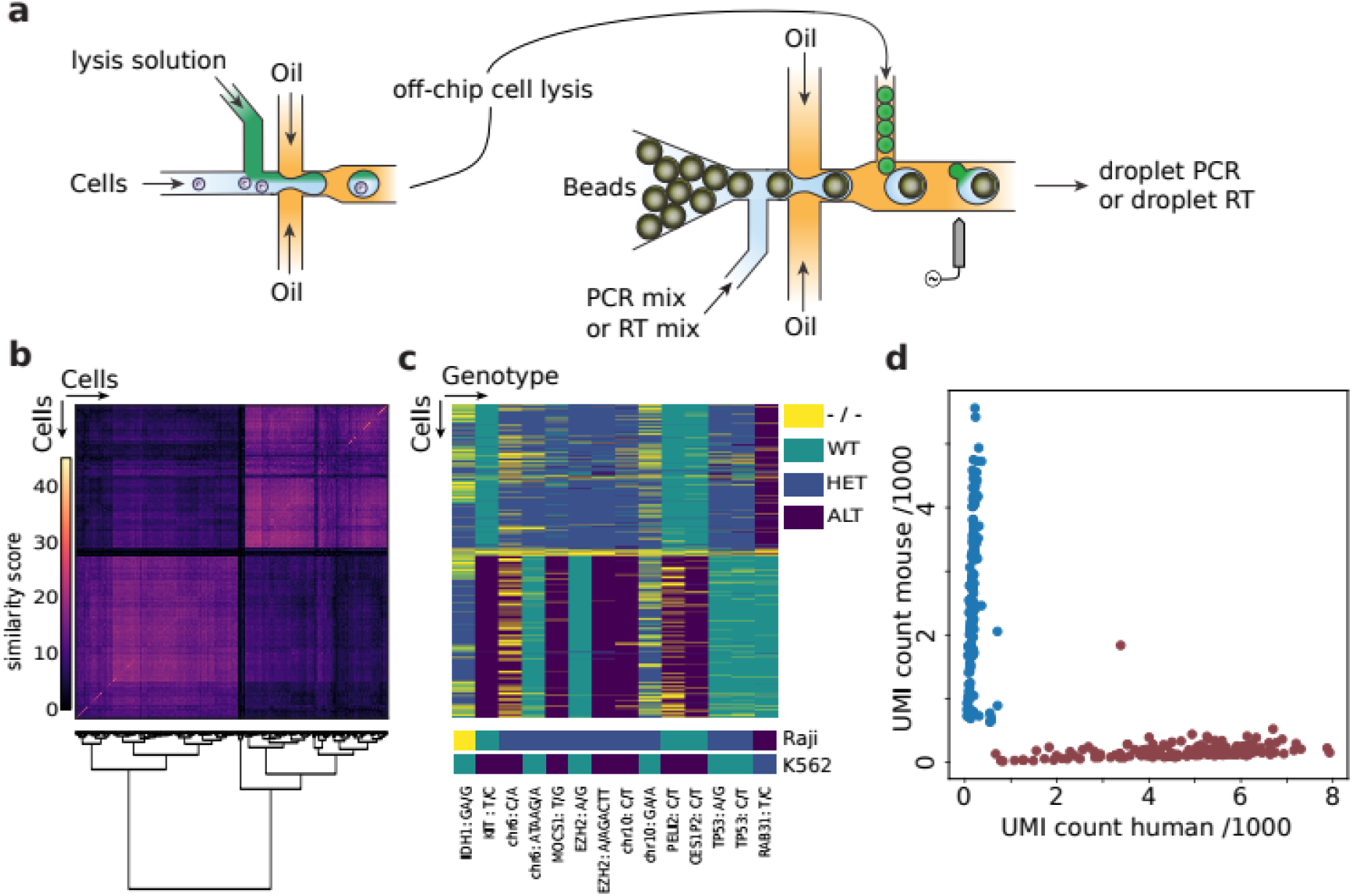
Singl-cell genotyping with barcode beads: **a** Schematic of the employed microfluidic two step protocol. A cell suspension is co-flowed with porteinase K or detergents and reinjected into a second device after off-chip incubation and heat inactivation of the protease. In the second device droplets with barcode beads are paired with cell lysate and PCR or RT reagents to create barcoded amplicons of targeted genomic regions. **b** Cell-cell similarity matrix based on the number of shared SNPs is given. Cells are ordered along both axes by hierarchical clustering using Ward’s minimum variance method. **c** Cells by genotype matrix, the cells (rows) are in the same order as in **b** and gnomic variants which are different between the two cell lines are given as columns. Genotyping dropouts are indicated in yellow. Bottom rows shows genomic variants detected from homogeneous bulk samples (full variant indexes are given in supplementary data table). **d** Single cell RNA sequencing result of the Mouse (3T3) Human (K562) species mixing experiment, counts represent detected unique molecular identifiers in thousands.

A common quality control experiment in single cell RNA sequencing is to profile a cell suspension of two species because individual transcripts can be assigned to cell type based on their sequence. To showcase the potential of the modular bead design for scRNAseq we therefore attach a poly-T primer instead of the cancer hot spot panel to the same bead batch and prepare a mixed mouse (NIH 3T3) and human (K562) cell sample. We use the same microfluidic workflow, but include an acidic lysis solution to stabilize RNA, and reverse transcription (RT) instead of PCR reagents, to produce a scRNAseq library. We collect droplets for ~1 minute aiming to capture 300-400 cells; sequencing yields 164 mouse cells with a median count of 2688 transcripts, 189 human cells with 5330 transcripts (Figure 2d). This demonstrates that our repurposed beads perform comparably to dedicated beads for scRNAseq (Supplementary figure 5a-f)(*2*). Bead modularity thus enables rapid deployment of the same barcode beads for single cell workflows targeting distinct molecules.

## Discussion

Our modular barcode design reduces the cost of bead synthesis while allowing rapid and inexpensive repurposing to other targets, including DNA and RNA. This makes them attractive for multiomic experiments by allowing fine tuning of primer sequences and concentrations to obtain best conditions for simultaneous DNA and RNA sequencing. This is important, because research on designing primers for multiplexed PCR reaction demonstrates that countering primer bias is a non trivial problem which requires empirical optimization to achieve good results(*28–31*). Moreover, the barcode sequences are compact and efficiently synthesized to full length, reducing sequencing waste and minimally consuming read length. A compact barcode structure also allows to use PCR primers to amplify specific cells of interest to increase their sequence coverage and improve signal quality in the pooled library(*32*). The ease with which highly multiplexed primers can be added enables new opportunities to enhance scRNAseq sensitivity. For example, poly-T mRNA capture yields an unbiased profile of expressed transcripts, but because coverage is typically below ~10%, low abundance transcripts may be missed. Our modular design might be used to enhance detection of such transcripts by dedicating a fraction of all primers on the bead to their capture alongside poly-T probes. Besides established DNA or RNA protocols, we expect that bead modularity will accelerate the introduction of novel assays by reducing the risk of fabricating dedicated bead batches that may not work.

## Methods

### Microfluidic device fabrication

Devices were fabricated with standard photolithography techniques(*33*). Custom device fabrication is not necessary to use these beads and can be substituted with commercially available instruments (e.g. from 10x Genomics, Mission Bio, 1CellBio and others). Master structures were made with Su8 3025 photoresist (MicroChem, Westborough, MA, USA) on a three inch silicon waver (University Wafer) by spin coating, soft baking at 95°C for 20 min and subjecting to 3 min UV-exposure through printed photolithography masks (CAD/Art Services, 12000 DPI)(supplementary file 2). Post UV exposure, the wafer was baked at 95°C for 2 min and cooled to room temperature and developed in a propylene glycol monomethyl ether acetate (Sigma Aldrich) bath, rinsed with acetate, and dried and hard baked at 225°C for 10 min. Curing agent and PDMS were mixed in 1:10 ratio, degassed and poured over the master structure and baked at 65°C for 4 h, removed from the master and punched with a 0.75 mm biopsy core (World Precision Instruments). The device was then bonded to a glass slide using O2 plasma and the channels were treated with Aquapel (PPG Industries) to render them hydrophobic. Aquapel was purged from the channels with air after 5 min contact time and residual liquid evaporated by baking at 65°C for 15 min.

### Barcode bead synthesis

Barcode beads were prepared by generating droplets on a microfluidic drop maker (supplementary figure 1) with an acrylamide premix (6% w/v Acrylamide, 0.15% w/v N, N’-Methylenebisacrylamide, 48 mM Tris-HCl pH 8.0, 0.3% w/v ammonium persulfate, 0.1x Tris-buffered saline–EDTA (TBSET: 10 mM Tris-HCL pH 8.0, 137 mM NaCl, 20 mM EDTA, 1.4 mM KCl, 0.1% v/v Triton-X100), 20 mM primer pBB1 (supplementary data table 7): /5Acryd/ACTAACAATAAGCTCUAUCGATGACCTAATACGACTCACTATAGGGACAAATGCC GATTCCTGCTGAAC (IDT) as dispersed phase and HFE-7500 (3M Novec) with 2% (w/v) PEG-PFPE amphiphilic block copolymer surfactant (008-Fluoro-surfactant, Ran Technologies) and 0.4% v/ v Tetramethylethylenediamine as continuous phase. The resulting emulsion was kept at room temperature over night to let the acrylamide polymerize. The emulsion was broken with 1H,1H,2H,2H-Perfluoro-1-octanol (Sigma Aldrich) and the beads washed three times in TBEST, three times in Tris-EDTA-Tween buffer (TET: 10 mM Tris-HCl pH 8.0, 10 mM EDTA, 0.1% v/v Tween-20), and three times in pre-ligation buffer (30 mM NaCl, 10 mM Tris-HCl pH 8.0, 1 mM MgCl2, 0.1% Tween-20). Beads were resuspended in T4 ligation buffer (NEB) and 7.5 μM of primer pBB2: /5Phos/TGACGTTCAGCAGGAATCGGCATTTGTCCCTATAGT GAGTCGTATTAGGTCATCGATAGAG at approximately 1:1 solid:solvent fraction. The suspension was heated to 75°C and slowly cooled to room temperature to anneal the primer pair.

Barcode primer plate and splint plate (supplementary data tables 1-6) for first round of split-pool barcode synthesis was prepared by combining each barcode with its cognate splint at a 1:1 ratio and a final concentration of 100 μM. Paired barcode and splints were phosphorylated in a PCR plate by combining 20 μl of oligonucleotides with 40 μl phosphorylation mix (1X T4 ligation buffer, 0.2 mg/ml BSA, 0.167 U/μl T4 PNK (NEB)) and incubated for 30 min at 37°C and heat inactivated for 20 min at 65°C. After phosphorylation, 100 μl of bead suspension to each barcode splint pair. To start the first round of ligation 40 μl of ligation mix (9.54 U/μl T4 ligase (NEB) in 1x T4 ligase buffer) was added to each well and the plate sealed and incubated for 1-4 h at room temperature and the enzyme inactivated by heating to 65°C for 10 min. Beads were collected, washed five times in TET and resuspended in ligation buffer at a 1:1 solid:solvent fraction for the next round of ligation. This process was repeated for the barcode fragments 2 and 3 to yield the final modular gel beads which can be quickly functionalized for a specific purpose.

To prepare the barcoded beads for single cell experiments we resuspended the beads at 1:2 solid to liquid in T4 ligation buffer and performed a splinted DNA ligation for 1 h at room temperature. For RNA beads we used 10 μM pBB4: CTCGAATAGG as splint and 10 μM pBB5: /5Phos/TTCGAGNNNNNNNNTTTTTTTTTTTTTTTTTTTTTTTTV as primer. For the cancer hot spot panel we phosphorylated a set of 49 primers (Mission Bio Acute Myeloid Leukemia Panel) or 16 primers (supplementary table) at equimolar ratio and ligated to the beads at 10 μM total concentration using 10 μM pBB8: CTGCGAGTACTAGG or pBB4 as splint. To render the barcodes single stranded, beads were washed four times in denaturing solution (100 mM NaOH, 0.5 % v/v Brij-35) and washing solution quenched by resuspending in low salt buffer (10 mM NaCl, 10 mM Tris-HCl pH 8.0, 0.1 mM EDTA, 0.1% Tween-20).

A detailed step-by-step description is provided as supplementary protocol.

### Barcode release test

Barcode beads (2 μl) were resuspended in LS (10 mM Tris-HCl pH 7.5, 1 mM MgCl2, 50 mM NaCl, 0.1% Tween20) and incubated for 1 minute. The beads were pelleted at 2000g, the supernatant removed and the beads resusbended in 20 μl 1x CutSmart (New England Biolabs), 1x Maxima H-RT buffer (Thermo Scientific) or 1x Kappa HiFi PCR buffer (Kapa Biosystems). For locally doublestranded cleavage tests pBB3 was added to a final concentration of 3 μM. Then, 0.4 μl USER II (New England Biolabs) enzyme mix was added an the suspension incubated for 45 min at 37°C, and heat inactivated. FAM-probe pBB6 was added to a final concentration of 1 μM and the beads incubate at room temperature for 15 minutes under rotation. Beads were washed 3 times in 1 ml TET and imaged on a fluorescence microscope (EVOS cell imaging systems, Thermo Fisher) at constant light intensity, shutter speed and signal amplification.

For bioanalyzer traces 1 μl beads were resuspended in 20 μl CutSmart and released with 0.5 μl USER II by incubating 30 minutes at 37°C. The Samples were diluted 4 fold with water and 1 μl supernatant was loaded on a Bioanalyzer High Sensitivity DNA electrophoresis chip (Agilent Technologies).

### Cell Culture

K562 (ATCC CCL-243); Raji (ATCC CCL-86), CCRF-CEM (ATCC CCL-119) and P493-6 (Cellosaurus CVCL_6783); NIH 3T3 (ATCC CRL-1658); or LAX7R(*34*) (gift from Jim Wells laboratory) cells were cultured at 37°C in the presence of 5% CO2 in Iscove’s Modified Dulbecco’s Medium; RPMI-1640 medium; Dulbecco’s Modified Eagle’s Medium; or MEMα with L-glutamine & Ribo-& Deoxyribonucleosides. Each media was supplemented with antibiotics and 10% fetal bovine serum (FBS). Human cell lines were washed in PBS, 3T3 were washed in PBS and detached by trypsinization to form a cell suspension for microfluidic experiments

### Cell encapsulation and barcoding

For the cancer hot spot experiment K562 and Raji cells or P493-6 and LAX7R cells were mixed at a 1:1 ratio and resuspended in phosphate-buffered saline (PBS) with 10.2% (w/v) Iodixanol at an approximate concentration of 3 million cells/ml. Cells were co-flowed with lysis buffer (100 mM Tris at pH 8.0, 0.5% IGEPAL, proteinase K 1.0 mg/mL)(*35*) each at 1500 μl/h and with 3000 μl/h HFE-7500 with 2% (w/v) PEG-PFPE amphiphilic block copolymer on a bubble-trigger device (*36*) (Supplementary file) to form droplets of about 45 μm diameter. Droplets were collected and incubated at 50°C for 1 h and 80°C 10 min to lyse the cells and heat inactivate proteinase K. On a bead-and-droplet merging device (Supplementary figure 1) closed packed, hot spot panel modified barcoding beads in DNA bead buffer (10 mM Tris-HCl pH 7.5, 40 mM NaCl, 2.5 mM MgCl2, 3.75% (v/v) Tween-20, 2.5% (v/v) Glycerol, 0.625 mg/ml BSA, 3 μM pBB3: TCATCGATAGAGCTTATTGT/3C6/) were reinjected at 75 μl/h and co-flowed with 150 μl/h PCR mix (1.65x NEBNext Ultra II Q5 Master Mix, 0.033 U/μl USER II (NEB), 1.32 M Propylene glycol, 0.25 mg/ml BSA, 0.5 mM DTT) to form bead containing droplets which were merged with the cell lysate containing droplets reinjected at 35 μl/h (about 200 Hz) by a salt water electrode(*37*) (Figure 2a). The droplets equivalent of about 1000 cells (69 s fractions) were collected, the oil excess oil removed from the collection tubes and replaced with FC40 with 5% (w/v) PEG-PFPE amphiphilic block copolymer. Droplet PCR was thermocycled with the following conditions: 30min at 30°C to release primers, 3 min at 95°C; 20 cycles of 20 s at 98°C, 10 s at 72°C, 4 min at 62°C, and 30 s at 72°C; and a final step of 2 min at 72°C with all ramp rates set to 1°C/s. Emulsion was broken with 1H,1H,2H,2H-Perfluoro-1-octanol diluted with 60 μl water the beads pelleted and 50 μl supernatant removed. To supernatant 5 μl 10x CutSmart (NEB) was added and incubated with 20 units ExoI nuclease (NEB) for 1 h at 37°C before purification with 42 μl AMPure beads (Beckman Coulter). The sample was eluted in 20 μl water which served as input for sequencing library generation. Cancer hotspot experiments with CCRF-CEM and K562 cells were done as previously described(*27*).

For the RNA experiment K562 and 3T3 cells were mixed at a 1:1 ratio and resuspended at 2.57 million cells/ml in the same buffer as above. Cells were co-flowed with lysis buffer (30 mM Na-citrate pH 6.5, 0.2 % Trition-X100, 0.2 % SDS, 2 mM EDTA, 10 mM DTT) on the same device as above, droplets collected and incubate 1 h at 4°C. Same bead-and-droplet merging device was used to combine pBB5 functionalized, closed packed barcoding beads in RNA bead buffer (1x Maxima H-RT Buffer (ThermoFisher), 2% (v/v) Tween-20, 0.625 mg/ml BSA, 3 μM pBB3). Beads were reinjected at 75 μl/h and co-flowed with 150 μl/h RT mix (1.65x Maxima H-RT Buffer, 0.033 U/μl USER II (NEB), 15 U/μl Maxima H-RT (ThermoFisher), 1.65 U/μl RNasin (Promega), 1.65 mM dNTP (NEB), 2.2 % (v/ v) Tween-20) and merged with the cell lysate containing droplets reinjected at 25 μl/h (about 200 Hz) same as above. The droplets equivalent of about 300-400 cells (60 s fractions) were collected and the oil exchanged as above. Emulsion was incubated at 37°C for 30 min then 54°C for 1 h. Emulsion was overlaid with 20 μl 1x CutSmart containing 20 units of ExoI, broken with 5 μl 1H,1H,2H,2H-Perfluoro-1-octanol, and incubated for 30 min at 37°C. RNA/DNA hybrids were purified with 40 μl AMPure beads and eluted in 20 μl water and stored at −20°C.

### Library preparation and sequencing

To the cancer hot spot library P5 and P7 sequences were attached by PCR using the custom pBB9 primer and Nextera N701 (Illumina). Library was purified with 0.6x volume fraction AMPure beads, its concentration measured by a fluorometer (Qubit 3.0, Invitrogen) and the absence of primer dimers verified on a Bioanalyzer High Sensitivity DNA electrophoresis chip (Agilent Technologies). For sequencing a MiSeq V2 300 cycles kit (Illumina) was used and the library diluted to 12 pM according to the recommendations of the sequencing kit manual. The library was sequenced in paired end mode and each side sequenced over 150 base pairs.

For RNA library, second strand synthesis and linear amplification by *in vitro* transcription (IVT) was done as described previously(*17*) without fragmenting RNA after IVT. IVT product was reverse transcribed with the primer: AAGCAGTGGTATCAACGCAGAGTGTANNNGGNNNB(*38*) as described(*17*), purified wit 0.9x volume fraction AMPure beads and the concentration measured by the Qubid dsDNA HS assay (Thermofisher). 13.8 ng RNA/DNA hybrid was fragmented with 20 μg Tn5(*39, 40*) (assembled with pBB11 and pBB12) in 50 μl. Library was amplified with pBB9 and pBB13 and NEBNext Ultra II Q5 according to the manufacturer protocol, purified with 0.65x AMPure beads and prepared a 12 pM sequencing library. The library was sequenced on a MiSeq V3 150 cycles kit (Illumina) in paired end mode, distributed 55/110 cycles, using custom sequencing primers pBB10 and pBB11 (supplementary data table 7).

### Bioinformatic data evaluation

Bead barcodes were parsed from the read 1 file with a custom script. The program cutadapt (v2.4) was used to find the common ligation scar between the combinatorial barcodes and the forward read primers. This step is necessary because the 0 – 3 base pair spacer in the barcode bead sequence (which helps cluster identification on the sequencing device by increasing sequence diversity for otherwise identical segments of the barcode sequence such as the ligation scars) prevents extraction of the combinatorial barcode blocks by distance from the 5’-end. Barcode blocks are extracted by distance from the position of the ligation scar (3’-side distance) and matched against a white list, allowing a maximum edit distance of one, to identify true barcode sequences.

Read 1 and read 2 sequences were demultiplexed into barcode groups and valid cell barcode groups were discriminated from background barcode groups by identifying the inflection point of the barcode rank plot versus number of associated reads. Reads from valid cell barcodes were processed as previously described(*12*). Briefly, FASTQ files with valid reads were aligned to the hg19 build of the human genome reference using bowtie2 (v2.3.4.1), filtered (properly mapped, mapping quality > 2, primary alignment), sorted, and indexed with samtools (v1.8). HaplotypeCaller from the GATK suite (v.4.1.3.0) was used to produce GVCF files and genotyped jointly on all genomic intervals with GATK GenotypeGVCFs. Genotyped intervals were combined into a single variant call format (VCF) file and multiallelic records split and left-aligned using bcftools (v1.9). Finally, variant records were exported to HDF5 format using a condensed representation of the genotyping calls (0: wildtype; 1: heterozygous alternate; 2: homozygous alternate; 3: no call). The result is a cell by allelic variant matrix, **V**, with the condensed genotype call as categorial matrix elements.

To cluster cells, genotype was one-hot encoded, converting **V** to four sparse binary matrices, **V_0_** to **V_3_**. Pairwise cell-cell similarity was calculated by computing the dot product for the matrices corresponding to heterozygous and alternate calls and summing them: **V_1_V_1_^T^ + V_2_V_2_^T^** yielding the cellcell similarity matrix **S**. Hierarchical clustering of **S** was done with SciPy (v1.3.1) using scipy.cluster.hierarchy.linkage and Ward’s minimum variance method. Variant calls that distinguish the two cell lines were identified by discarding all calls for which less than 10% of cells were called as alternate (sum of heterozygous and homozygous alternate) yielding 27 potentially informative variants. Remaining variant calls were inspected and discarded if constant over all cells. The remaining 14 variant call positions are the final list.

RNA sequencing data was evaluated with Kallisto(*41*) and Bustool(*42*). Since Kallisto currently does not support variable length barcode sequences, the 0 – 3 base pair spacer was deleted from each read by a custom script. Next, Kallisto bus with option -x 0,0,24:0,24,32:1,0,0 was run to extract barcode sequences and pseudo align transcripts. Skipping barcode correction by the whitelist approach, output was converted to gene counts with Bustool and visualized in python using Scanpy(*43*) and matplotlib(*44*).

## Supporting information

supplementary figures and protocol

## Acknowledgments

We thank Sarah Pyle for help with the figure art and members from the Abate laboratory for valuable discussions.

## Author Contributions

**CLD**: Conceptualization, Methodology, Software, Formal analysis, Investigation, Writing – Original Draft **ARA**: Supervision, Funding acquisition, Writing – Review & Editing.

## Data and materials availability

All scripts are available on GitHub at https://github.com/AbateLab/ModularGelBeads. All sequencing data generated in this study is available on the Sequence Read Archive under BioProject number PRJNA632423 and PRJNA660010 upon final publication.

## Supplementary Materials

Supplementary file 1: Bead fabrication protocol, supplementary figures and table Supplementary data tables: Primer sequences, amplicon positions and variant calls Supplementary file 2: microfluidic devices

## References and Notes

1. Shapiro, E., Biezuner, T. and Linnarsson, S. (2013) Single-cell sequencing-based technologies will revolutionize whole-organism science. Nat. Rev. Genet., 14, 618–630.

2. Klein, A.M., Mazutis, L., Akartuna, I., Tallapragada, N., Veres, A., Li, V., Peshkin, L., Weitz, D.A. and Kirschner, M.W. (2015) Droplet Barcoding for Single-Cell Transcriptomics Applied to Embryonic Stem Cells. Cell, 161, 1187–1201.

3. Macosko, E.Z., Basu, A., Satija, R., Nemesh, J., Shekhar, K., Goldman, M., Tirosh, I., Bialas, A.R., Kamitaki, N., Martersteck, E.M., et al. (2015) Highly Parallel Genome-wide Expression Profiling of Individual Cells Using Nanoliter Droplets. Cell, 161, 1202–1214.

4. Abate, A.R., Chen, C.-H., Agresti, J.J. and Weitz, D.A. (2009) Beating Poisson encapsulation statistics using close-packed ordering. Lab Chip, 9, 2628.

5. Shalem, O., Sanjana, N.E., Hartenian, E., Shi, X., Scott, D.A., Mikkelsen, T.S., Heckl, D., Ebert, B.L., Root, D.E., Doench, J.G., et al. (2014) Genome-Scale CRISPR-Cas9 Knockout Screening in Human Cells. Science, 343, 84–87.

6. Parnas, O., Jovanovic, M., Eisenhaure, T.M., Herbst, R.H., Dixit, A., Ye, C.J., Przybylski, D., Platt, R.J., Tirosh, I., Sanjana, N.E., et al. (2015) A Genome-wide CRISPR Screen in Primary Immune Cells to Dissect Regulatory Networks. Cell, 162, 675–686.

7. Pellegrino, M., Sciambi, A., Treusch, S., Durruthy-Durruthy, R., Gokhale, K., Jacob, J., Chen, T.X., Geis, J.A., Oldham, W., Matthews, J., et al. (2018) High-throughput single-cell DNA sequencing of acute myeloid leukemia tumors with droplet microfluidics. Genome Res., 28, 1345–1352.

8. Shahi, P., Kim, S.C., Haliburton, J.R., Gartner, Z.J. and Abate, A.R. (2017) Abseq: Ultrahigh-throughput single cell protein profiling with droplet microfluidic barcoding. Sci. Rep., 7, 44447.

9. Stoeckius, M., Hafemeister, C., Stephenson, W., Houck-Loomis, B., Chattopadhyay, P.K., Swerdlow, H., Satija, R. and Smibert, P. (2017) Simultaneous epitope and transcriptome measurement in single cells. Nat. Methods, 10.1038/nmeth.4380.

10. Peterson, V.M., Zhang, K.X., Kumar, N., Wong, J., Li, L., Wilson, D.C., Moore, R., McClanahan, T.K., Sadekova, S. and Klappenbach, J.A. (2017) Multiplexed quantification of proteins and transcripts in single cells. Nat. Biotechnol., 35, 936–939.

11. Chen, S., Lake, B.B. and Zhang, K. (2019) High-throughput sequencing of the transcriptome and chromatin accessibility in the same cell. Nat. Biotechnol., 10.1038/s41587-019-0290-0.

12. Demaree, B., Delley, C.L., Vasudevan, H.N., Peretz, C.A.C., Ruff, D., Smith, C.C. and Abate, A.R. (2020) Joint profiling of proteins and DNA in single cells reveals extensive proteogenomic decoupling in leukemia. bioRxiv, 10.1101/2020.02.26.967133.

13. Dixit, A., Parnas, O., Li, B., Weissman, J.S., Friedman, N., Dixit, A., Parnas, O., Li, B., Chen, J., Fulco, C.P., et al. (2016) Perturb-Seq: Dissecting Molecular Circuits with Scalable Single-Cell RNA Profiling of Pooled Genetic Resource Perturb-Seq: Dissecting Molecular Circuits with Scalable Single-Cell RNA Profiling of Pooled Genetic Screens. Cell, 167, 1853-1857.e17.

14. Jaitin, D.A., Weiner, A., Yofe, I., Lara-Astiaso, D., Keren-Shaul, H., David, E., Salame, T.M., Tanay, A., van Oudenaarden, A. and Amit, I. (2016) Dissecting Immune Circuits by Linking CRISPR-Pooled Screens with Single-Cell RNA-Seq. Cell, 167, 1883–1896.e15.

15. Adamson, B., Norman, T.M., Jost, M., Parnas, O., Regev, A. and Weissman, J.S. (2016) A Multiplexed Single-Cell CRISPR Screening Platform Enables Systematic Dissection of the Unfolded Protein Response. Cell, 167, 1867–1882.

16. Mazutis, L., Gilbert, J., Ung, W.L., Weitz, D.A., Griffiths, A.D. and Heyman, J.A. (2013) Single-cell analysis and sorting using droplet-based microfluidics. Nat. Protoc., 8, 54–56.

17. Zilionis, R., Nainys, J., Veres, A., Savova, V., Zemmour, D., Klein, A.M. and Mazutis, L. (2016) Single cell barcoding and sequencing using droplet microfluidics. Nat. Protoc., 12, 44–73.

18. Levenshtein, V.I. (1966) Binary Codes Capable of Correcting Deletions, Insertions and Reversals. Sov. Phys. Dokl., 10, 707.

19. Faircloth, B.C. and Glenn, T.C. (2012) Not all sequence tags are created equal: Designing and validating sequence identification tags robust to indels. PLoS One, 7.

20. Horspool, D.R., Coope, R.J.N. and Holt, R.A. (2010) Efficient assembly of very short oligonucleotides using T4 DNA Ligase. BMC Res. Notes, 3.

21. Zhang, F., Christiansen, L., Thomas, J., Pokholok, D., Jackson, R., Morrell, N., Zhao, Y., Wiley, M., Welch, E., Jaeger, E., et al. (2017) Haplotype phasing of whole human genomes using bead-based barcode partitioning in a single tube. Nat. Biotechnol., 35, 852–857.

22. Saikia, M., Burnham, P., Keshavjee, S.H., Wang, M.F.Z., Heyang, M., Moral-Lopez, P., Hinchman, M.M., Danko, C.G., Parker, J.S.L. and De Vlaminck, I. (2019) Simultaneous multiplexed amplicon sequencing and transcriptome profiling in single cells. Nat. Methods, 16, 59–62.

23. Grosselin, K., Durand, A., Marsolier, J., Poitou, A., Marangoni, E., Nemati, F., Dahmani, A., Lameiras, S., Reyal, F., Frenoy, O., et al. (2019) High-throughput single-cell ChIP-seq identifies heterogeneity of chromatin states in breast cancer. Nat. Genet., 51,1060–1066.

24. Gérard, A., Woolfe, A., Mottet, G., Reichen, M., Castrillon, C., Menrath, V., Ellouze, S., Poitou, A., Doineau, R., Briseno-Roa, L., et al. (2020) High-throughput single-cell activity-based screening and sequencing of antibodies using droplet microfluidics. Nat. Biotechnol., 10.1038/s41587-020-0466-7.

25. Wang, Y., Cao, T., Ko, J., Shen, Y., Zong, W., Sheng, K., Cao, W., Sun, S., Cai, L., Zhou, Y., et al. (2020) Dissolvable Polyacrylamide Beads for High-Throughput Droplet DNA Barcoding. Adv. Sci., 10.1002/advs.201903463.

26. Ward, J.H. (1963) Hierarchical Grouping to Optimize an Objective Function. J. Am. Stat. Assoc., 58, 236–244.

27. Delley, C.L. and Abate, A.R. (2020) Microfluidic particle zipper enables controlled loading of droplets with distinct particle types. Lab Chip, 10.1039/D0LC00339E.

28. Markoulatos, P., Siafakas, N. and Moncany, M. (2002) Multiplex polymerase chain reaction: A practical approach. J. Clin. Lab. Anal., 16, 47–51.

29. Shen, Z., Qu, W., Wang, W., Lu, Y., Wu, Y., Li, Z., Hang, X., Wang, X., Zhao, D. and Zhang, C. (2010) MPprimer: a program for reliable multiplex PCR primer design. BMC Bioinformatics, 11, 143.

30. Sint, D., Raso, L. and Traugott, M. (2012) Advances in multiplex PCR: balancing primer efficiencies and improving detection success. Methods Ecol. Evol., 3, 898–905.

31. Carlson, C.S., Emerson, R.O., Sherwood, A.M., Desmarais, C., Chung, M.-W., Parsons, J.M., Steen, M.S., LaMadrid-Herrmannsfeldt, M.A., Williamson, D.W., Livingston, R.J., et al. (2013) Using synthetic templates to design an unbiased multiplex PCR assay. Nat. Commun., 4, 2680.

32. Ranu, N., Villani, A.-C., Hacohen, N. and Blainey, P.C. (2019) Targeting individual cells by barcode in pooled sequence libraries. Nucleic Acids Res., 47, e4–e4.

33. Qin, D., Xia, Y. and Whitesides, G.M. (2010) Soft lithography for micro- and nanoscale patterning. Nat. Protoc., 5, 491–502.

34. Pollock, S.B., Hu, A., Mou, Y., Martinko, A.J., Julien, O., Hornsby, M., Ploder, L., Adams, J.J., Geng, H., Markus, M., et al. (2018) Highly multiplexed and quantitative cell-surface protein profiling using genetically barcoded antibodies. 10.1073/pnas.1721899115.

35. Eastburn, D.J., Sciambi, A. and Abate, A.R. (2013) Ultrahigh-Throughput Mammalian Single-Cell Reverse-Transcriptase Polymerase Chain Reaction in Microfluidic Drops. Anal. Chem., 85, 8016–8021.

36. Yan, Z., Clark, I.C. and Abate, A.R. (2017) Rapid Encapsulation of Cell and Polymer Solutions with Bubble-Triggered Droplet Generation. Macromol. Chem. Phys., 218, 1600297.

37. Sciambi, A. and Abate, A.R. (2014) Generating electric fields in PDMS microfluidic devices with salt water electrodes. Lab Chip, 14, 2605–2609.

38. Hughes, T.K., Wadsworth, M.H., Gierahn, T.M., Do, T., Weiss, D., Andrade, P.R., Ma, F., Silva, B.J. de A., Shao, S., Tsoi, L.C., et al. (2019) Highly Efficient, Massively-Parallel Single-Cell RNA-Seq Reveals Cellular States and Molecular Features of Human Skin Pathology. bioRxiv, 10.1101/689273.

39. Picelli, S., Björklund, A.K.A.K., Reinius, B., Sagasser, S., Winberg, G. and Sandberg, R. (2014) Tn5 transposase and tagmentation procedures for massively scaled sequencing projects. Genome Res., 24, 2033–2040.

40. Di, L., Fu, Y., Sun, Y., Li, J., Liu, L., Yao, J., Wang, G., Wu, Y., Lao, K., Lee, R.W., et al. (2020) RNA sequencing by direct tagmentation of RNA/DNA hybrids. Proc. Natl. Acad. Sci. U. S. A., 117, 2886–2893.

41. Bray, N.L., Pimentel, H., Melsted, P. and Pachter, L. (2016) Near-optimal probabilistic RNA-seq quantification. Nat. Biotechnol., 34, 525–527.

42. Melsted, P., Ntranos, V. and Pachter, L. (2019) The barcode, UMI, set format and BUStools. Bioinformatics, 10.1093/bioinformatics/btz279.

43. Wolf, F.A., Angerer, P. and Theis, F.J. (2018) SCANPY: large-scale single-cell gene expression data analysis. Genome Biol., 19, 15.

44. Hunter, J.D. (2007) Matplotlib: A 2D Graphics Environment. Comput. Sci. Eng., 9, 90–95.

