## supplementary figures and protocol for "Modular barcode beads for microfluidic single cell genomics"

### Supplementary Tables and Figures: Modular barcoding beads for microfluidic single cell genomics

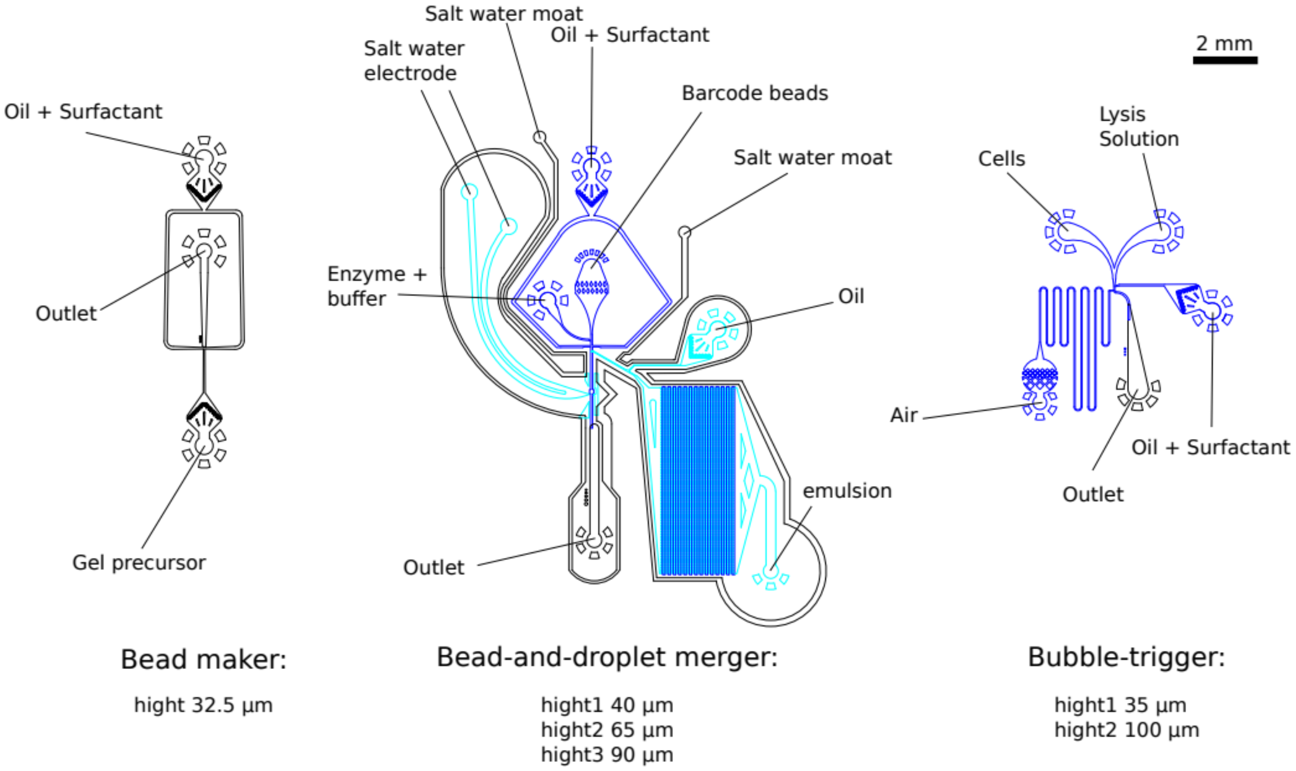


**Supplementary figure 1, devices:** Overview over the devices used in this manuscript (see also supplementary file 2).

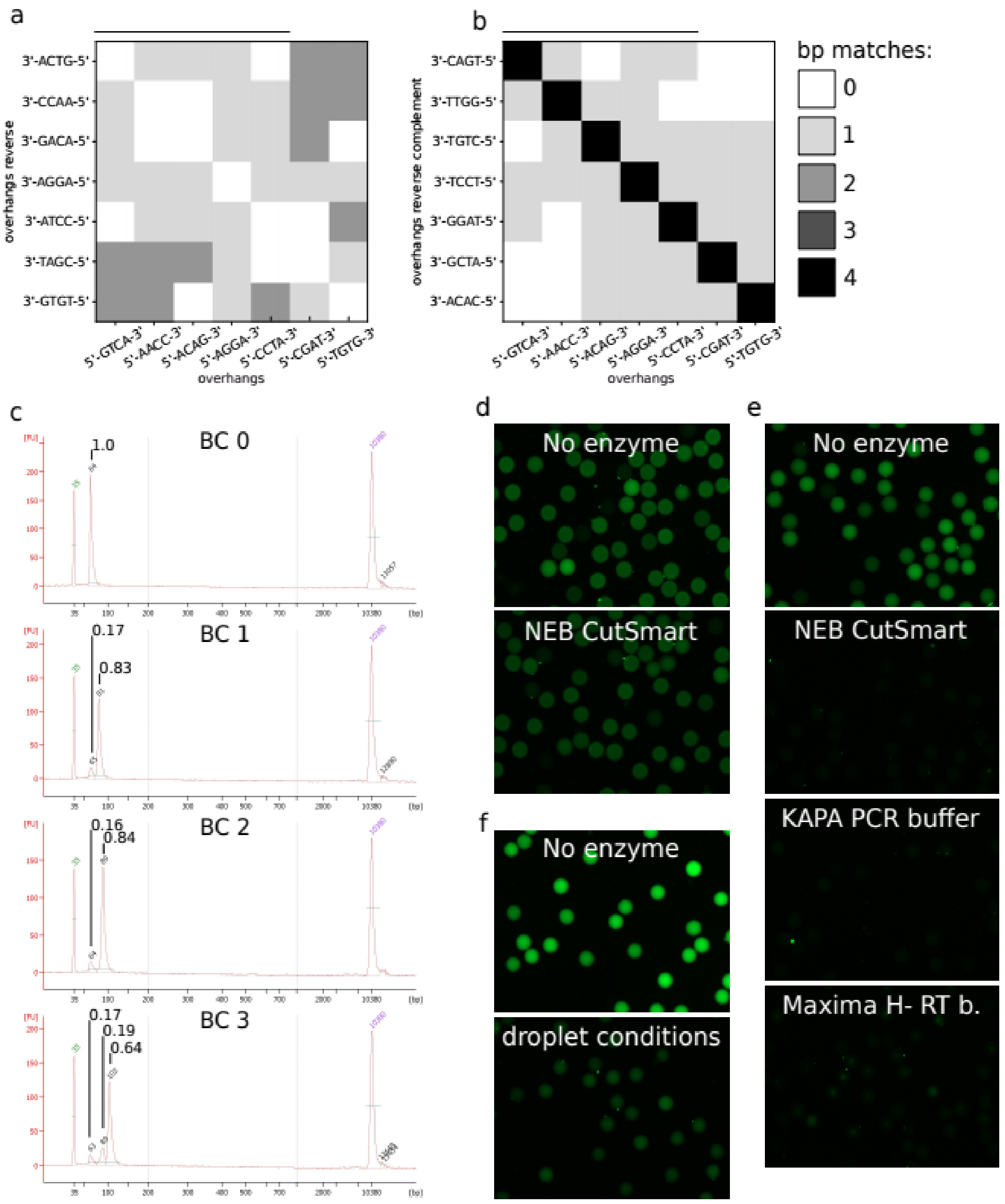


**Supplementary figure 2, Split-pool ligation of barcode beads: a** Matrix of number of base pair matches between different ligation handles and self complementary. Low matching reduces potential for undesired barcode products and concatamer formation. **b** Same as **a** for the reverse complement of the ligation handles. Black bar above columns in **a** and **b** indicates employed handles in this article. **c** Barcode intermediates (top 3) and full length (bottom) released from the beads after the indicated barcode fragment ligation (0-3) were analyzed with a bioanalyzer. Small peak numbers indicate fragment size and large peak number molar fraction of molecules in the respective peak. **d, e, f** FAM-labeled reverse complement probe of the PCR handle was hybridized to the barcode beads after completing fabrication and imaged with a fluorescence microscope. Disappearance of fluorescence is measured after incubation for 45 minutes at 37°C with and without USER II enzyme mix and different buffers. **d** Single stranded barcodes are no good substrate. **e** Barcodes are efficiently released after hybridization of a 20-bp complementary probe (pBB3) in all tested buffers including reverse transcription and PCR buffer, and **f** in droplet conditions of the single-cell genotyping experiment. Bead diameter is about 55 um.

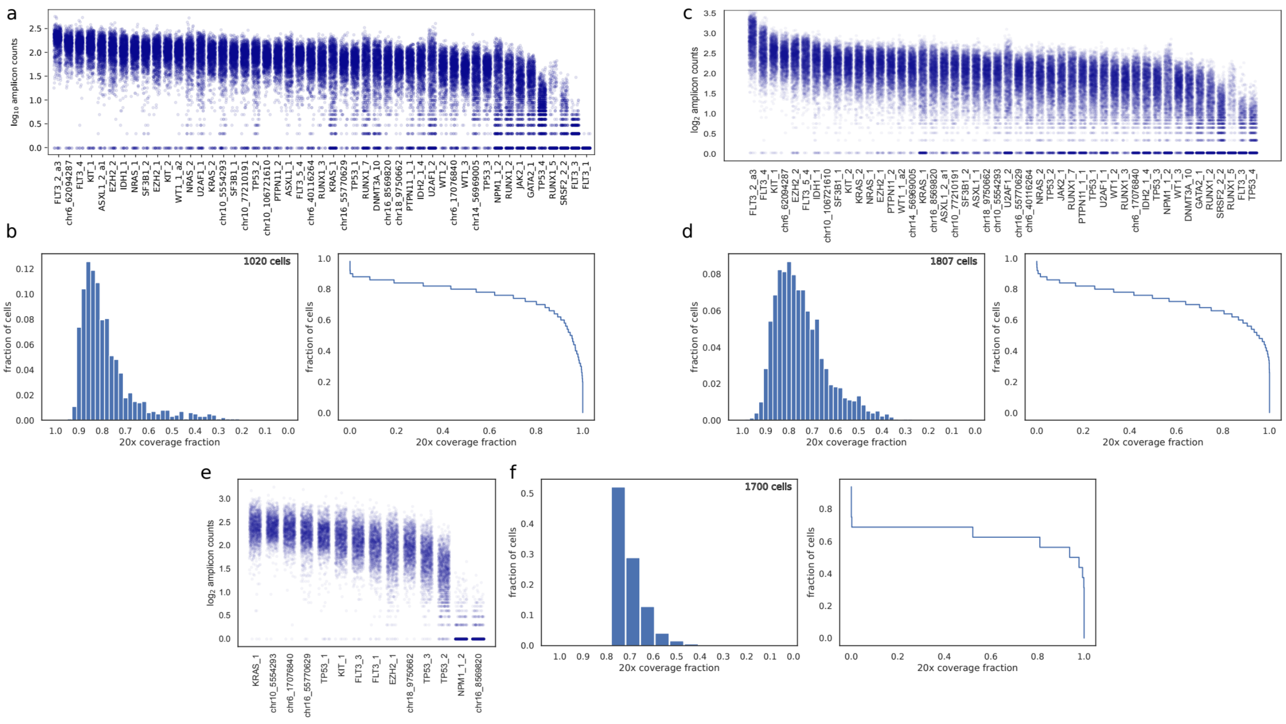


**Supplementary figure 3, Primer dropout ad panel uniformity for single cell genomics experiments: a** Amplicon count per cell and target genomic loci for the Raji and K562 experiment (figure 2b and 2c). For each target genome locus the number of observed amplicon reads per cell is depicted as log 10 counts. Some genomic locations are harder to amplify by single cell PCR resulting in more amplicon dropout (e.g. FLT3 amplicon 1 and 3). **b** histogram (left) and cumulative plot (right) of the fractions of targeted amplicons with at least 20 fold coverage per cell demonstrating that about 70% of cells cover 80% of the targeted amplicons with at least 20 reads. **c** and **d** same as **a** and **b** for the P493-6 and LAX7R experiment (supplementary figure 5a). **e** and **f** same as **a** and **b** for the K562 and CEM experiment (supplementary figure 5b)

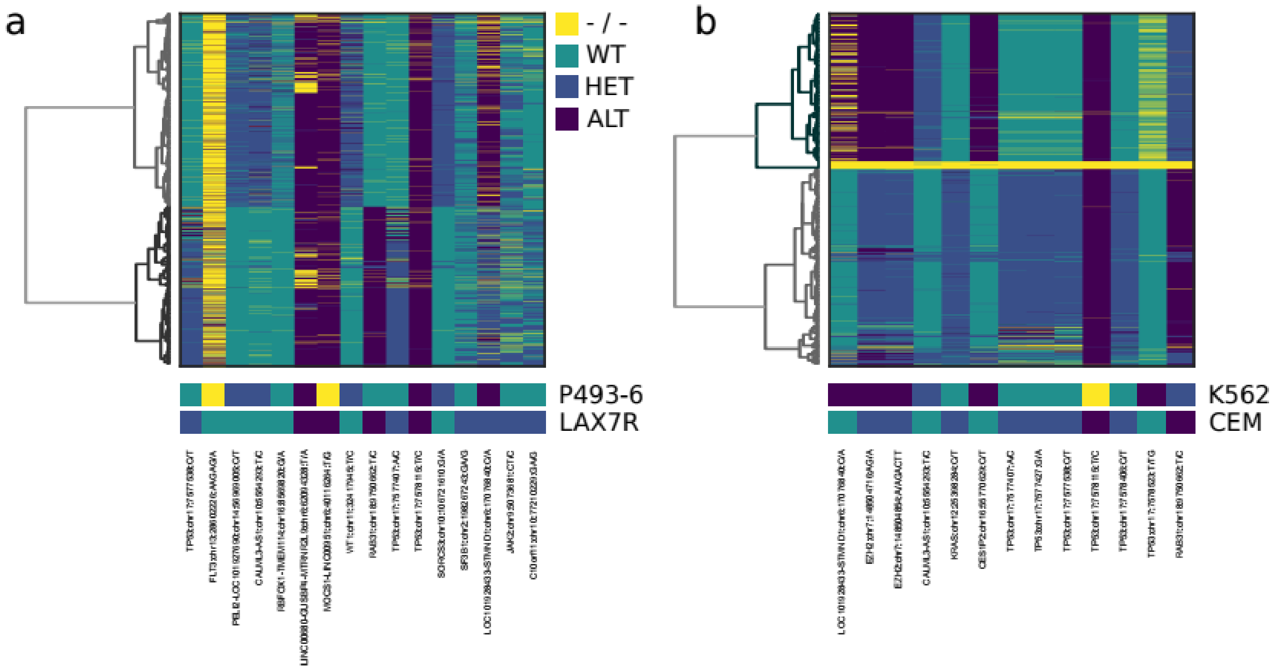


**Supplementary figure 4, single cell genotype profiling for different cell lines: a** Single cell genotype matrix (cells by genotype) as in figure 2c for P493-6 and LAX7R cells. Pairwise cell similarity was defined as sum of shared alternate calls over all target loci and clustered by Ward's minimum variance method (see methods). A total of 1809 cells were profiled. **b** Same as a for 1620 K562 and CEM cells, which were profiled with a 16 primer set (a subset of the primers in a) for K562 and CEM cells.

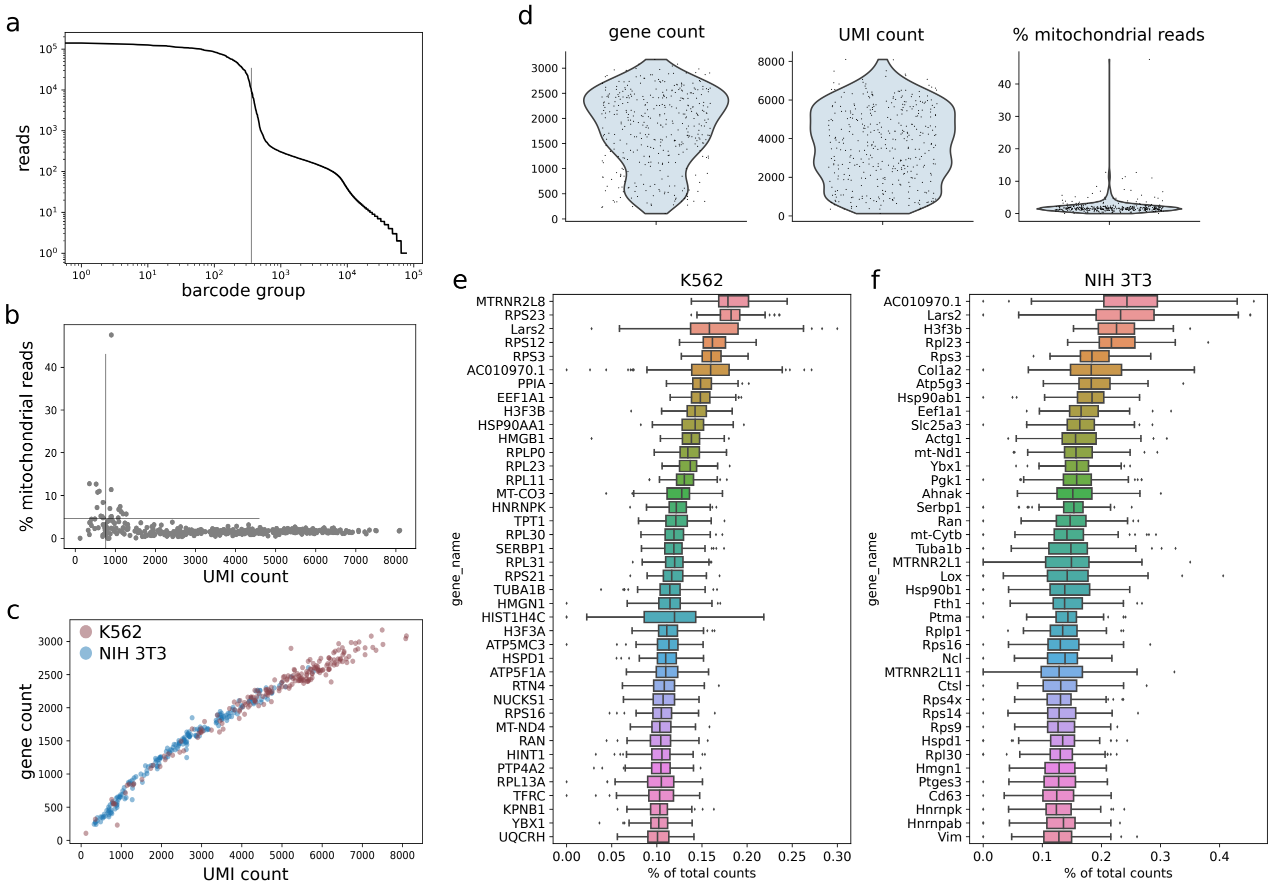


**Supplementary figure 5, single cell RNA-seq data quality: a** Barcode rank versus reads plot exhibiting "knee" shape. b UMI counts versus percent mitochondrial reads. Barcode groups were required to feature at least 775 UMI counts and less than 5% mitochondrial reads to be considered a cell, which yields 345 cells. c UMI counts versus gen counts demonstrates that the data set has not been sequenced to saturation. Scatters are colored according to inferred cell type demonstrating a bias in transcript number between the two cell types in line with previous observations. d Violin plots of common quality metrics: gen count per cell, UMI count per cell and percent mitochondrial reads per cell. e and f 40 detected most highly expressed genes for K562 and 3T3 cells, respectively. A majority code for ribosomal, mitochondrial or histone components which is in line with metabolically active and dividing cells.

**Supplementary Table 1: Bead fabrication cost overview**: Bead costs of InDrops beads are compared to the new approach. Costs are calculated based on list prices of vendors and are given as "set-up cost" that is the up front investment to start bead fabrication and as "cost per ml" reflecting the steady state cost per batch.

**InDrops Gel Beads** **746496 barcode space**

| | | scale order | yield guarantee | | set-up cost | | bead volume yield | | cost per ml bead | | cost per experiment (~ 20 000 cells) | | source |
|---|---|---|---|---|---|---|---|---|---|---|---|---|---|
| 1 | InDrops Acrydite-modified p. | 10000 | 125 | nmol | 3218.8 | $ | 5.7 | ml | 569.7 | $ / ml | 28.48 | $ | IDT |
| 18 | Barcode Plates | 10 | - | nmol | 15042.6 | $ | 80.0 | ml | 188.0 | $ / ml | 9.40 | $ | IDT |
| 1 | BST 2.0 | 8000 | - | units | 283.0 | $ | 8.6 | ml | 33.0 | $ / ml | 1.65 | $ | NEB |
| 1 | dNTP | 40 | - | umol | 250.0 | $ | 14.8 | ml | 16.9 | $ / ml | 0.85 | $ | NEB |
| | **total** | | | | **18794.4** | $ | | | **807.7** | $ / ml | **40.38** | $ | |

(125 nmol is guaranteed yield, calculated cost for 250 nmol because IDT usually delivers more than the minimum)

**Ligation Gel Beads** **884736 barcode space**

| | | scale order | yield guarantee | | set-up cost | | bead volume yield | | cost per ml bead | | cost per experiment (~ 20 000 cells) | | source |
|---|---|---|---|---|---|---|---|---|---|---|---|---|---|
| 1 | pBB1 | 10000 | 1600 | nmol | 1738.4 | $ | 72.3 | ml | 24.0 | $ / ml | 1.20 | $ | IDT |
| 3 | Barcode Plates | 25 | - | nmol | 902.9 | $ | 12.0 | ml | 75.2 | $ / ml | 3.76 | $ | IDT |
| 3 | Splint Plates | 100 | - | nmol | 1199.5 | $ | 50.0 | ml | 24.0 | $ / ml | 1.20 | $ | IDT |
| 1 | pBB2 (phos-lig_start) | 10000 | 1600 | nmol | 1107.2 | $ | 72.3 | ml | 15.3 | $ / ml | 0.77 | $ | IDT |
| 1 | pBB3 (release-3C) | 1000 | 100 | nmol | 102.0 | $ | 16.0 | ml | 6.4 | $ / ml | 0.32 | $ | IDT |
| 1 | pBB4 end splint | 1000 | 200 | nmol | 21.0 | $ | 10.7 | ml | 2.0 | $ / ml | 0.10 | $ | IDT |
| 1 | pBB5 (phos-RNA) | 1000 | 200 | nmol | 126.9 | $ | 10.7 | ml | 11.9 | $ / ml | 0.59 | $ | IDT |
| 1 | T4 PNK | 2500 | - | units | 228.0 | $ | 13.0 | ml | 17.5 | $ / ml | 0.88 | $ | NEB |
| 1 | T4 Ligase | 100000 | - | units | 256.0 | $ | 4.2 | ml | 61.4 | $ / ml | 3.07 | $ | NEB |
| 1 | USER | 250 | - | units | 297.0 | $ | 8.3 | ml | 35.6 | $ / ml | 1.78 | $ | NEB |
| | **total** | | | | **5978.9** | $ | | | **273.4** | $ / ml | **13.67** | $ | |

(1600 nmol is guaranteed yield, calculated cost for 3200 nmol because IDT usually delivers more than the minimum)

### Supplementary Protocol for Barcode Bead Fabrication: Modular barcoding beads for microfluidic single cell genomics

Cyrille L. Delley and Adam R. Abate

We order our primers typically from IDT, all enzymes and biochemical reagents from NEB and the chemicals from Sigma. Buffer and solution formulations and chemicals are attached at the end of this document and primer tables are given as a separate document. The description is for a bead batch of 5 ml but can easily be scaled to other volumes. A typical microfluidic single cell experiment aiming for 20,000 cells consumes about 50 µl beads. The protocol can be completed in two full days.

**Order barcode fragment plates and prepare for experiments**

1| We typically order the barcode fragment plates and splints separately because splints require a larger fabrication scale due to their short length. Order all primer plates with primers resuspended in water at 200 µM and normalized yield to the maximum available. Add an equal volume of each splint from splint plate 1 to the corresponding well of barcode plate 1 with a multi pipette and repeat for the other barcode and splint plates. All barcode plates are now at 100 µM and have double stranded barcode fragments. Reseal the plates with a fresh aluminum foil and take care not to cross contaminate the wells. Store at -20 °C.

2| Make your dropmaker in PDMS according to previous protocols.

note 1: instead of making your own dropmaker a variety of comercial vendors are available. Look for a simple dropmaker with one inlet for the aqueous polyacrylaminde solution and one inlet for the fluorinated oil. Depending on your target gel bead size, aim for droplets in the range 35 - 60 µm in diameter. After breaking of the emulsion we typically observe a gel bead diameter / droplet diameter expansion of about 1.13 fold. A 53 µm droplet will yield a 60 µm gel.

note 2: smaller droplets tend to polymerize less reliably than larger ones. Droplets of 40 µm diameter and larger rarely fail in our hands.

**Bead Fabrication**, (2.5 h hands-on, overnight incubation)

3| Prepare six 1.5 ml collection tubes by filling them with 300 µl mineral oil. Label the first one with “waste” and the others with tube 1 to 5

4| Prepare 3.4 ml of HFE-7500 with 2% surfactant and 0.4% v/v TEMED, mix well, load into a 5 ml syringe, prime and connect to microfluidic device. (Oil volume depends on the flow rates used and might require adjustments for different drop makers)

5| Prepare the following solution:

| Component | stock | final | | Volume /ul |
|---:|:---:|:---:|:---:|:---:|
| Acrylamide 40%/5% Bis | 40.00 | 3.00 | w/v % | 187.50 |
| contains Bisacrylamide | 5.00 | 0.150 | w/v % | - |
| Acrylamide 40% | 40.00 | 3.00 | w/v % | 187.50 |
| APS | 30.00 | 0.30 | w/v % | 25.00 |
| pBB1 | 100.00 | 20.00 | uM | 500.00 |
| TBSET | 1.00 | 0.10 | x | 250.00 |
| Tris-HCl pH 8.0 | 1000 | 48 | mM | 120.00 |
| H2O | - | - | - | 1230 |
| **total** | | | | **2500** |

note 3: pBB1 is not light sensitive and no special precautions against premature cleavage have to be taken

<u>note 4: Monomeric acrylamide is toxic and carcinogenic, use appropriate precautions</u>

6| Fill a 3 ml syringe with 300 µl HFE-7500 and overlay with well mixed acrylamide solution from point 7. Prime the syringe, mount in a vertical position pointing upwards and connect to microfluidic device. Connect a short tubing to the device outlet and place into the “waste” tube. This tube will collect the polydispersed outflow prior to the stabilization of the flows and will be discarded after the run.

7| For our device, set the flow rate to 1000 µl/h for the acrylamide mix and 1250 µl/h for the oil. For other devices, tune the flow rates to make the desired droplet size. We typically aim for 45 - 55 µm droplet diameter. Wait about 1 min or until flows stabilize and move outlet to the first collection tube. Move outlet to the next tube once they fill up, about every 30 min.

8| Once a tube is full, close the lid and move to an incubator set to 60°C. Tubes should be incubated 2 hours after witch polymerization is usually complete but can go overnight. Overnight incubation at room temperature (RT) works equally well unless when dealing with small amounts of emulsion.

**Bead cleanup**, (2 h hands-on)

9| Centrifuge the collection tubes at 100 g, 30 s and use a 200 µl gel loading tip to remove the HFE oil from the bottom. Multiple aspirations will be needed. There is no need to replace the tip between aspirations or different tubes. Overlay the beads with 1 ml TBSET and centrifuge at 3000 g, 30 s. Remove leftover HFE from each tube and remove mineral oil overlay together with TBSET by pipetting or aspiration. Take care not to aspirate the bead emulsion. Repeat TBSET step once more.

9| Add 800 µl TBSET and 150 µl of 20% (v/v) 1H,1H,2H,2H-Perfluoro-1-octanol in HFE 7500 (20% PFO) to each tube, briefly vortex at full power and place on overhead rotator for 5 min. Spin down at 3000 g, 30 s and remove the 20% PFO mix with a gel loading tip from the bottom then the mineral oil emulsion and TBSET from the top without removing the beads.

10| Repeat step 9 but reduce centrifugal force to 500 g and skip removal of 20% PFO from the bottom. After removing TBEST from the top, replace with 1 ml fresh TBEST and gently kick up the bead pellet by pipetting up and down without emulsifying the 20% PFO. Place a 70 µm strainer on a 50 ml tube and combine the beads from all tubes by transferring the bead emulsions to the strainer. Flush the strainer with TBSET until only bead precipitates remain on top.

11| Centrifuge tube at 2000 g for 3 min, remove the supernatant, resuspend the pellet with 10 ml TET and transfer to a 15 ml tube. Wash beads 3 times by spinning down at 2000 g for 2 min and resuspending with fresh 10 ml of TET.

Stopping point: beads can be kept at 4°C for 1 year at this point.

12| Optionally, repeat bead fabrication and clean up (step 3 - 11) to prepare a larger batch for barcoding.

**Split-pool protocol, 1st round,** (1 h hands on, 2 h 15 min total)

The following is calculated for 5 ml bead volume. For other volumes scale accordingly and make sure the PCR plate can hold the total volume

13| Thaw the barcode plate 1 with the added splints

14| Once thawed, phosphorylate primers with T4 polynucleotide kinase (PNK). Prepare the enzyme master mix:

| | | | | | |
|---|---|---|---|---|---|
| T4 lig buffer | 10 | 1.00 | x | 600 | ul |
| H2O | - | - | - | 3550 | ul |
| BSA | 20 | 0.06 | mg/ml | 18 | ul |
| T4 PNK | 10 | 0.05 | U/ul | 32 | ul |

and add 40 µl master mix to each well of a PCR plate. Then add 20 µl from each well of the barcode plate 1 to the corresponding well of the PCR plate with a multi pipette. Seal the plate with aluminum foil and incubate for 30 min at 37°C and heat inactivate 20 min at 65°C

15| While primers are phosphorylating, wash beads 3 times with 10 ml PL buffer. Resuspend the beads according to the table and heat to 75°C for 2 min and let slowly cool down to RT to anneal the primer pBB2:

| | | | | | |
|---|---|---|---|---|---|
| Beads | - | - | - | 5000 | ul |
| T4 lig buffer | 10 | 1 | x | 1000 | ul |
| pBB2 | 100 | 3.75 | uM | 750 | ul |
| H2O | - | - | - | 3250 | ul |

Once primer phosphorylation is complete remove the foil and distribute 100 µl beads to each well. Store a 5 µl bead aliquot in a PCR tube, labeled "round 0" for quality control.

9| Prepare the T4 ligase master mix and distribute 40 µl to each well and mix by pipetting up and down. We use a multi pipette and a trough. Make sure to replace the tips after each pipette cycle to not cross contaminate the wells.

| | | | | | |
|---|---|---|---|---|---|
| T4 lig buffer | 10 | 1 | x | 420 | ul |
| H2O | - | - | - | 3760 | ul |
| T4 ligase | 2000 | 1.91 | U/ul | 20 | ul |

Seal the plate and incubate at RT for 1 h (longer incubation works too). Heat inactivate at 65°C for 10 min and cool to RT.

16| Use a multi pipette and collect the well contents into a trough filled with 15 ml TET. There is no need to replace the tips in this step. Distribute collected bead suspension equally between two 15 ml tubes. Centrifugate at 2000 g for 3 min, remove the supernatant. Wash five times in 15 ml TET. The beads are now ready for the next split-pool cycle. You can stop at any time during the washes and proceed at another day. In this case leave the beads suspended in TET.

**Split-pool protocol, 2$^{nd}$ and 3$^{rd}$ round** (each 1 h hands on, 2 h 15 min total)

17| Proceed as for the first round using barcode plate 2 or barcode plate 3. There is no need to add more pBB2 to the beads in point 8| since the barcode fragments are already double stranded and you can replace the pBB2 volume with water. Always at step 15 store a 5 µl bead aliquot in a PCR tube, labeled with the corresponding cycle number for quality control. Take an additional QC sample after the last round of split-pool ligation.

18| After the 3$^{rd}$ split-pool round the beads are fully barcoded but without functional primer ends. This is our preferred stopping point, because any primer end can now be ligated quickly enabling flexible bead usage in different genomic experiments (e.g. for scRNA-seq we ligate a poly-T primer with UMI or for targeted genomics a panel of primers against the sites of interest). Beads suspended in TET at 4°C are stable for at least 12 months.

**Primer functionalization and bead cleanup** (1 h hands on, 2 h total)

19| Take the appropriate amount of beads from your prefabricated bead stock (50 µl corresponds to about one experiment and we rarely functionalize less than 250 µl beads per round). The following example is calculated for 1000 µl pellet volume and is done in a 15 ml tube (use smaller tubes for bead volumes below 500 µl). Wash the beads 3 times with 10 ml PL buffer and remove supernatant. Add the follwig reagents without the enzyme:

| | | | | | |
|---|---|---|---|---|---|
| T4 lig buffer | 10 | 1.00 | x | 300.00 | ul |
| primer end mix | 100 | 10.00 | uM | 300.00 | ul |
| pBB4 | 100 | 10.00 | uM | 300.00 | ul |
| Beads | - | - | - | 1000.00 | ul |
| H2O | - | - | - | 1095.05 | ul |
| T4 ligase | 2000 | 3.3 | U/ul | 4.95 | ul |
| **Total** | | | | **3000** | **ul** |

Mix well by vortexing, incubate for 5 min at RT, then add the enzyme and mix by inverting the tube 20 times and place it on a overhead rotator for 1 h at RT.

note 5: primer end mix corresponds to the mixture of targeting primers which should be linked to the beads. E.g. for a single cell RNA seq experiment we use pBB5 or for the cancer hot spot experiment we use an equimolar mixture of 50 cancer hot spot forward primers (each primer at 0.2 µM). Primers need to be 5' phosphorylated.

20| Remove a 5 µl aliquot for QC and pellet beads at 2000g for 3 min, remove supernatant. Wash once in 10 ml H2O and 4 times in 5 ml denaturing solution to make primer single stranded. Incubate beads for 3 min at RT and overhead rotation during washes. After last wash remove supernatant, wash 3 times in 10 ml LS buffer to quench the base.

21| Optionally perform bead cleanup to remove incompletely fabricated bead primers. To clean the beads, add 2 volumes of 1x NEB buffer 3.1 to 1 volume of bead pellet. Add 3.5 µM (final solution concentration) of complementary primers to your targeting ends. Head beads to 75°C and let slowly cool to RT. Add 1 µl of ExoI nuclease per 50 µl of bead pellet and incubate for 1 h at RT. Heat inactivate ExoI and repeat step 20 to remove the complementary primers from the beads

22| Either store beads in LS until first use or resuspend in your running buffer now. Add an equimolar amount of pBB3 to aid in USER mediated barcode release. The on bead primer concentration of the bead pellet is about 5 µM for beads after cleanup (20 µM starting concentration times 0.5 sphere packing density times 0.5 primer fabrication efficiency) and 10 µM for beads without cleanup (incomplete primers also present a binding site)

**Quality control by FAM-probes**

23| Resuspend the beads, take 5 µl aliquots and transfer to 1.5 ml spin tubes. Add 25 µl TET to each tube, add 1 ul of pBB6 (100 µM stock) to the first tube and 1 µl pBB7 to the second tube. Vortex and incubate 5 min at RT. Wash 3 times with 1 ml of TET and image beads under a fluorescence microscope. Always use the same illumination parameters for your quality control images to keep them comparable. Beads should light-up homogeneously as in supplementary figure 1d-f.

**Quality control by Bioanalyzer**

24| Prepare 30 µl 2x NEB Cut Smart and add 1.25 µl of NEB USERII enzyme. To each QC tube (round 0 - 3 and post functionalization) add 5 µl USER mix, mix by pipetting and incubate 30 min at 37°C. Add 40 µl water to each tube, mix and pellet beads. Load 1 µl of each tube to a Bioanalyzer DNA high sensitivity lane. See supplementary figure 1c how a successful fabrication should look.

**Buffers and Solutions**

| | | Stock | Final | Volume /ml |
|---|---|---|---|---|
| Tris-buffered saline–EDTA –Triton buffer (TBSET) | H2O | - | - | 821.7 |
| | Tris-HCL pH 8.0 | 1 M | 10 mM | 10.0 |
| | NaCl | 1 M | 137 mM | 137.0 |
| | EDTA pH 8.0 | 0.5 M | 10 mM | 20.0 |
| | Trito X-100 | 10 % (v/v) | 0.10 % (v/v) | 10.0 |
| | KCl | 2 M | 2.7 mM | 1.4 |
| | | | | **1000.0** |
| Denaturing solution | H2O | - | - | 20.8 |
| | NaOH | 1 N | 150.00 mN | 3.8 |
| | Brij-35 | 30 % (w/w) | 0.50 % (w/v) | 0.4 |
| | | | | **25.0** |
| Tris–EDTA–Tween buffer (TET) | H2O | - | - | 480.0 |
| | Tris-HCl pH 8.0 | 1 M | 10.00 mM | 5.0 |
| | EDTA pH 8.0 | 0.5 M | 10.00 mM | 10.0 |
| | Tween-20 | 10 % (v/v) | 0.10 % (v/v) | 5.0 |
| | | | | **500.0** |
| Low salt buffer (LS) | H2O | - | - | 488.9 |
| | NaCl | 5 M | 10.00 mM | 1.0 |
| | Tris-HCl pH 8.0 | 1 M | 10.00 mM | 5.0 |
| | EDTA pH 8.0 | 0.5 M | 0.10 mM | 0.1 |
| | Tween-20 | 10 % (v/v) | 0.10 % (v/v) | 5.0 |
| | | | | **500.0** |
| Pre ligation buffer (PL) | H2O | - | - | 486.5 |
| | NaCl | 5 M | 30 mM | 3.0 |
| | Tris-HCl pH 7.5 | 1 M | 10 mM | 5.0 |
| | Tween-20 | 10 % (v/v) | 0.1 % (v/v) | 5.0 |
| | MgCl2 | 1 M | 1 mM | 0.5 |
| | | | | **500.0** |

**Chemicals**

Primers (see supplementary table, IDT)
Acrylamide 40% (Sigma, A4058-100ML)
Acrylamide, Bisacrylamide 40%, 19:1 (Sigma, A9926-100ML)
N′-Tetramethyl ethylenediamine (Sigma, T9281-25ML)
Ammonium peroxydisulfate (Sigma, A3678-25G)
Brij-35 (Sigma, 203724-100ML)
Tween-20 (Sigma, P9416-100ML)
Trition X-100 (Sigma, T8787-100ML)
Tris(hydroxymethyl)aminomethane (Sigma, 252859-500G)
EDTA, 0.5 M, pH 8.0 (Sigma, 324506-100ML)
NaCl (Sigma, S9888-500G)
$MgCl_2$, 1 M (Sigma, M1028-100ML)
KCL (Sigma, P3911-500G)
BSA, Molecular Biology Grade (NEB, B9000S)
Thermolabile Exonuclease I (NEB, M0568L)
T4 DNA ligase (NEB, M0202M)
T4 Polynucleotide Kinase (NEB, M0201L)
Thermolabile USER II Enzyme (NEB, M5508L)
1H,1H,2H,2H-Perfluoro-1-octanol (Sigma, 370533-25G)
Novec HFE-75000 (3M)
PEG-PFPE amphiphilic block copolymer surfactant (008-Fluoro-surfactant, Ran Technologies)
Aquapel (PPG Industries)
